# MicroRNA Networks Driving Skeletal Aging and WNT Pathway Modulation

**DOI:** 10.64898/2026.01.23.701417

**Authors:** David Achudhan, David Monroe, Mrunal K. Dehankar, Ling Qin, Sundeep Khosla, Robert J. Pignolo, Abhishek Chandra

## Abstract

Cellular senescence is a key mechanism of skeletal aging in both physiological and accelerated conditions, such as radiotherapy. This study aimed to identify common differentially regulated microRNAs (miRs) across these contexts. We performed miR sequencing on three models: femurs from young (5-month-old) versus old (24-month-old) mice; focally radiated versus non-radiated femurs; and osteocytes from young versus old mice. Osteocytes were included in the comparison, as they have the longest lifespan in the mineralized bone matrix and they form 90-95% of all mesenchymal bone cell types. Among the three groups, miR-135a-5p and miR-671-5p were the common (i.e., shared) miRs that were downregulated, and miR-183-5p, a miR that regulates the WNT pathway, was the only shared upregulated miR, while miR-155-5p, a miR that regulates the Senescence-Associated Secretory Phenotype (SASP), was elevated in two conditions. The WNT-pathway has been positively associated with bone health and Sclerostin, a WNT-pathway inhibitor produced and secreted by osteocytes, has been implicated in accelerated skeletal deterioration following radiation. Thus, we used a neutralizing antibody to Sclerostin (Scl-Ab), to assess genes related to the WNT pathway and senescence, which are regulated by miR-183-5p and miR-155-5p, respectively. We further performed miR sequencing in radiated bones from mice treated with Scl-Ab and identified miR-133a-3p, a key miR that inhibits bone metabolism and function, which is upregulated in accelerated skeletal aging (i.e., focal radiation) downregulated by Scl-Ab. Overall, our study identifies potential regulatory gene pathways that modulate skeletal aging in the presence and absence of a WNT activator, Scl-Ab.

## Introduction

Cellular senescence has been identified as a key underlying mechanism for skeletal aging and several age-related morbidities. It is characterized by cells entering a state of stable survival, utilizing protective pathways to become resistant to signals that would normally trigger apoptosis^1^. The onset of cellular senescence is primarily induced by conditions that challenge the cell’s integrity and homeostasis. These conditions encompass damage (like DNA damage and genomic instability), metabolic strain (oxidative stress), and disruptions to the genome, specifically telomere shortening and compromised proteostasis^2^. With aging and disease conditions, the immune system also becomes less efficient at clearing senescent cells, which then persist. Senescent cells are metabolically active, with an elevated expression of pro-inflammatory proteins known as the Senescence-Associated Secretory Phenotype (SASP). Presence of senescence and the SASP have been well-established as a cause for aging-related frailty and multimorbidity^3–7^.

Osteoporosis, an age-related morbidity, is considered a key hallmark of bone aging, and is accompanied by a high risk for fractures^8^. Compared to men and younger women, post-menopausal women, following the loss of estrogen, are at an even higher risk of bone loss and subsequent fractures^9^. Some hallmarks of skeletal aging, such as loss of functional osteoblasts, increase in osteocyte apoptosis, empty lacunae, increase in bone marrow adiposity (BMAT) and an increase in the number and activity of osteoclasts, are seen in both older individuals of either sex, as well as in in post-menopausal women. However, certain hallmarks of skeletal aging, such as cellular senescence, the SASP, cell-fate switching in mesenchymal cells toward production of bone marrow adipocytes (BMAds) or reduction in osteoblast mineralization tend to be more prominent in age-related rather than post-menopausal osteoporosis.

Elevated expression of the genes related to cyclin-dependent kinase inhibitors, several SASP genes and genes associated with BMAd, have supported that cellular senescence plays a role in age- and radiation-associated skeletal aging and in defining several features of skeletal aging^10–12^. Genes that regulate features of skeletal aging are themselves controlled by microRNA (miRNA or miRs), short double-stranded RNA entities that bind to these genes to regulate their function.

To identify common miRNAs that can define skeletal aging from a regulatory perspective, we performed an unbiased comparison of miRNAs that are expressed in old bones and compared them with miRNAs expressed in focally radiated bones. Furthermore, since osteocytes compose a large proportion of bone cells, we also included miRNA sequencing data sets from old osteocytes. This three-way comparison is thus an attempt to narrow the list of miRNAs that can begin to define skeletal aging but are also unique to aged osteocytes. There are several cell-signaling pathways that regulate bone metabolism, but one of the key pathways that regulate bone metabolism is the WNT pathway. Previously, we have shown that WNT pathway activation was key in suppressing acute DNA damage and cellular apoptosis of bone cells following exposure to radiation^13,14^. Our study here further explores the relationship between skeletal aging-related miRNAs and WNT pathway regulation. This relationship is further explained using a neutralizing antibody (Scl-Ab) to Sclerostin, a WNT-pathway inhibitor protein, secreted mainly by the osteocytes.

## Methods

### Animal study design

All animal studies were approved by the Institutional Animal Care and Use Committee at the Mayo Clinic. The animals were purchased from The Jackson Laboratory and housed in our facility at 23–25°C with a 12-h light/dark cycle and were fed with standard laboratory pellets with free access to water. For the micro RNA sequencing from aged bones and osteocyte-enriched bone fraction (vertebrae treated with collagenase), was used as described earlier^10^ and the RNA was processed as described^15^.

The gene expression (by qRT-PCR) and miRNA sequencing data was generated using non-radiated (NR) and radiated (R) samples at 2 weeks post radiation, related to vehicle- and Scl-Ab-treated mice, as described previously^13^. Briefly, all animal procedures were reviewed and approved by the Institutional Animal Care and Use Committee (IACUC) at the University of Pennsylvania. Ten-week-old C57BL/6 mice were obtained from the Jackson Laboratory (Bar Harbor, ME, USA) and housed in groups under controlled conditions (23–25°C, 12-hour light/dark cycle) with free access to water and standard chow. Focal irradiation was performed on the distal metaphyseal region of the right femur using SARRP (Xstrahl, Suwanee, GA, USA) delivering a cumulative dose of 16Gy (8 Gy on days 1 and 3). Radiation was delivered to a 5 × 5 mm collimated field centered approximately 1 mm below the growth plate at 1.65 Gy/min, guided by built-in μCT and X-ray imaging. For Scl-Ab treatment, mice were randomized into two groups with comparable body weights: vehicle (isotonic buffer, Novartis) or Scl-Ab (100 mg/kg/week, Novartis) was administered once weekly by subcutaneous injections as described previously^13^, and samples were analyzed on day 14.

### RNA preparation and Quantitative RT-PCR

A 5 mm region below the growth plate of the distal metaphyseal femur was cut out from the radiated and the non-radiated legs, or young and old mice. After cleaning of any additional muscle tissue, the bone samples were stored in TRIzol (Invitrogen) at -80°C. Bones were homogenized and total RNA was isolated using a phase separation method, followed by a RNase free DNase treatment to remove genomic DNA and cleaning of samples with the RNeasy Mini Columns (QIAGEN, Valencia, CA). The quality of the RNA was determined using Nanodrop spectrophotometer (Thermo Scientific, Wilmington, DE). The cDNA was generated from mRNA using the High-Capacity cDNA Reverse Transcription Kit (Applied Biosystems by Life Technologies, Foster City, CA), according to the manufacturer instructions.

Primers which were earlier designed for our previous studies^10^ were used for rt-qPCR in this study as well. Additional primers were designed using Primer Express ® Software Version 3.0 (Applied Biosystems, Foster City, CA). Using a high throughput ABI Prism 7900HT Real Time System (Applied Biosystems, Carlsbad, CA) with SYBR Green (QIAGEN, Valencia, CA) as the detection method, gene expression was detected. The gene expression was normalized against the average of 2 reference genes (Actb and Tuba1) and the difference between the reference gene and the target gene (reference Ct- gene of interest Ct) or fold change was calculated comparing the median cycle threshold (Ct) from the radiated femurs with the non-radiated femurs as control.

### NEBNext miRNA Sequencing

Small RNA libraries were prepared using 500 ng of total RNA according to the manufacturer’s instructions for NEBNext Multiplex Small RNA Kit (New England Biolabs; Ipswich, MA). After purification of the amplified cDNA constructs, the concentration and size distribution of the PCR products was determined using an Agilent Bioanalyzer DNA 1000 chip (Santa Clara, CA) and Qubit fluorometry (Invitrogen, Carlsbad, CA). Four of the cDNA constructs were pooled and the 120-160bp miRNA library fraction was selected using Pippin Prep (Sage Science, Beverly, MA). The concentration and size distribution of the completed libraries were determined using an Agilent Bioanalyzer DNA 1000 chip (Santa Clara, CA) and Qubit fluorometry (Invitrogen, Carlsbad, CA).

Library pools (6 samples/lane) were loaded onto paired end flow cells at concentrations of 300 pM to generate cluster densities of 700,000/mm2 and ∼100 million reads per sample following Illumina’s standard protocol using the Illumina cBot and HiSeq 3000/4000 PE Cluster Kit. The flow cells were sequenced as 51 X 2 paired end reads using the Small RNA Sequencing Primer and an Index read to facilitate demultiplexing of the samples. Libraries were sequenced on an Illumina HiSeq 4000 using HiSeq 3000/4000 sequencing kit and analysis was performed using HCS v3.3.20 collection software. Base-calling is performed using Illumina’s RTA version 2.5.2.

### Secondary analysis

CAP-miRSeq version 1.2^16^ is a miRNASeq bioinformatics pipeline developed at the Mayo Clinic for comprehensive analysis of raw microRNA sequencing paired-end reads. The CAP-miRSeq workflow accepts unaligned reads (FASTQ) as input. The workflow generates raw and normalized expression counts for both known mature miRNAs and predicted novel miRNAs. Aligned reads (BAMs aligned to mm10), single nucleotide variants (SNVs), and various quality reports are also generated. The CAP-miRSeq workflow integrates following features to streamline data analysis: 1) read quality assessment and trimming using Cutadapt ^17^; 2) fast alignment through Bowtie ^18^ to different RNA species of genome; 3) mature/precursor/star miRNA detection and quantification using MiRDeep2 ^19^; 4) novel miRNA prediction and quantification using MiRDeep2 ^19^; 5) data visualization using IGV and 6) SNV detection in the seed region of miRNAs using GATK.

### Differential expression (DE) analysis

QC checks were performed on the cohort using principal component analysis, hierarchical clustering and visualization of raw and CPM normalized counts. Differential miRNA expression analysis was performed using R package edgeR. The criteria for selection of significant differentially expressed microRNAs were: log2 fold change > |1| and Benjamini-Hochberg adjusted p-value < 0.05. *x* significant differentially expressed microRNAs were identified using these thresholds.

### 4.5 Statistical analysis

All statistics were performed using GraphPad Prism 8.1.1 software. Data are expressed as median with interquartile range and analyzed by an unpaired two-tailed t-test when comparing two groups or a two-way ANOVA, when comparing two or more groups and variables. Heat maps were created using Morpheus software.

## Results

### Identification of shared miRNAs involved in skeletal aging

Gene expression of a single gene can be regulated by multiple miRs and a single miR can regulate multiple genes. To identify the miRs that play an important role in skeletal aging, miRNA-sequencing was performed in three pairs of samples: (i) femurs collected from young (5 month) and old (aged 24 month), (ii) an enriched osteocyte population from young (5 month) and old (aged 24 month) and (iii) a 5mm femur metaphyseal sample from non-radiated (4 month) and 16Gy focally radiated (4 month) femurs at 2 weeks post-radiation. 440 miRs were differentially expressed in whole femur from old (O) mice as compared to young (Y) mice, while 173 miRs were expressed in the enriched osteocytes from O bones when compared to bones from Y mice, while 96 miRs were differentially expressed in R-femurs as compared to the contralateral NR-femurs (Fig. 1A, B). The miRs that are upregulated or downregulated in all three conditions are shown using heat maps (Supp. Figs. 1-3). Venn diagrams were generated using either downregulated or upregulated miRs from O-femurs, R-femurs and O-osteocytes (Fig.1A, B). To identify common miRs that regulate either physiological or pathological skeletal aging, we focused on shared miRs which were at the intersection of all three conditions, O-femurs, R-femurs and O-osteocytes. While there were unique miRs that were downregulated in each of the three skeletal aging conditions, there were two in common miRs, miR-671-5p and miR-135a-5p, that were downregulated in all three conditions (Fig.1A). Surprisingly, miR-183-5p was the only upregulated miR in all three skeletal conditions (Fig. 1B), suggesting important pathway(s) that this single miR regulates.

**Figure 1.**
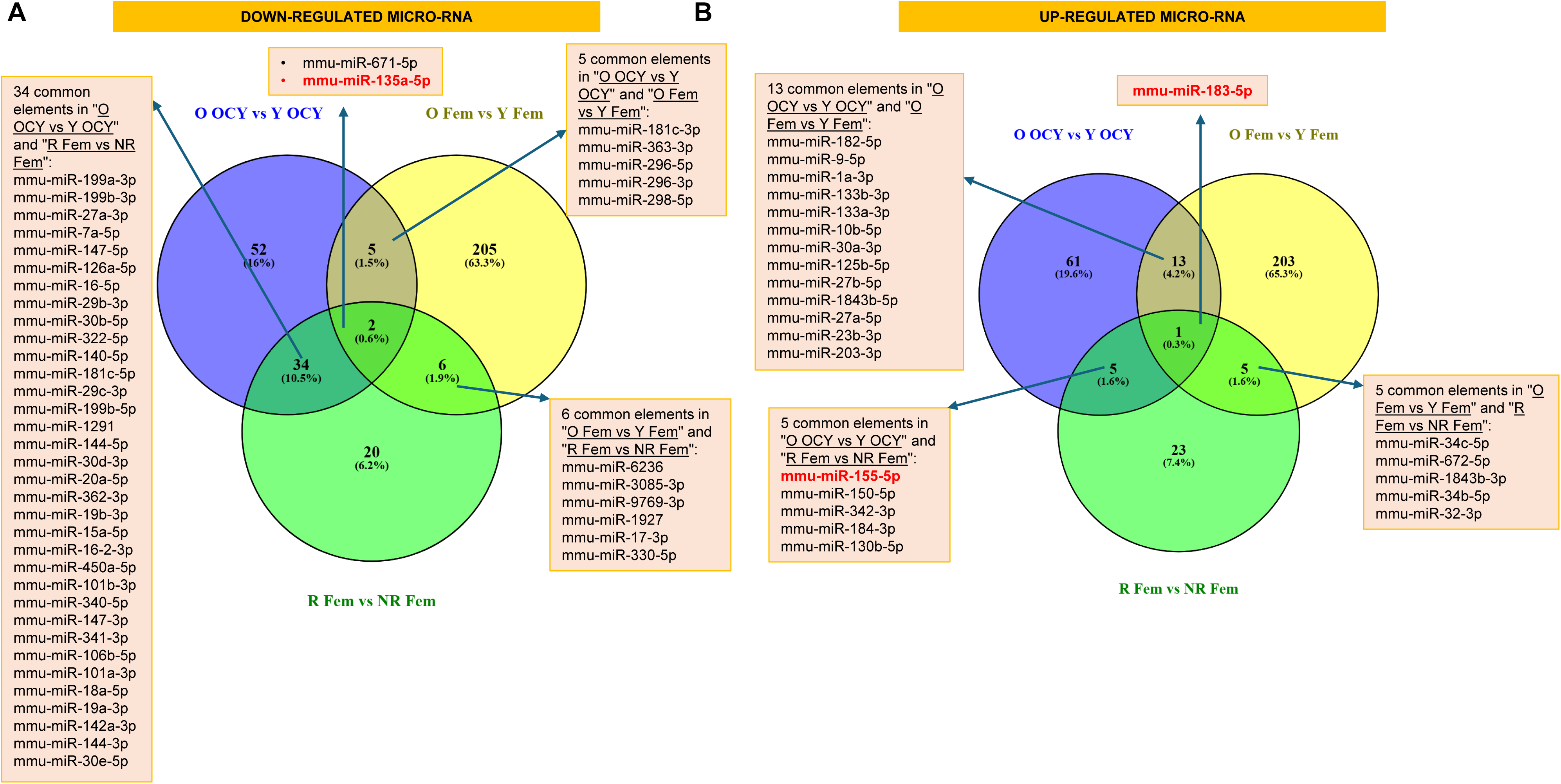
Comparative analysis of miRNAs expressed during skeletal aging. (A) MicroRNA (miR) sequencing was performed in RNA samples from O- vs Y- osteocyte enriched population, R- vs NR-femurs and O- vs Y- femurs. Figure represents the differentially expressed miRs in the O- vs Y- osteocyte enriched population, R- vs NR-femurs and O- vs Y-femurs. Venn diagram shows common or exclusive down-regulated miRs in O- vs Y- osteocyte enriched population, R- vs NR-femurs and O- vs Y- femurs. (B) Venn diagram shows common or exclusive up-regulated miRs in O- vs Y- osteocyte enriched population, R- vs NR-femurs and O- vs Y- femurs.

### WNT pathway gene regulation by miR-183-5p during skeletal aging

As mentioned above, miR-183-5p was the only upregulated miR in all three conditions of skeletal aging (O-femurs, R-femurs and O-osteocytes) (Fig.2A). Bioinformatics network analysis using miRTargetLink 2.0^20^, revealed that a major target of miR-183-5p was the WNT signaling pathway. Among the genes that are regulated by miR-183-5p, was Fzd1 (Frizzled Class Receptor 1), a G-protein-coupled receptor crucial for WNT signaling pathway (Fig. 2B). Interestingly, analysis of miR-sequencing data from vehicle- and Scl-Ab- treated mouse femurs revealed that miR-183-5p was significantly downregulated in Scl-Ab treated mouse femurs as compared to vehicle treated femurs (Fig. 2C). To test if elevation of miR-183-5p would result in the downregulation of WNT pathway, we analyzed samples collected from radiated bones and saw a significant decline in several WNT genes (as shown in the heat map, Fig.2D). In a separate study, in which animals were randomized to receive Scl-Ab, a WNT pathway activator, we could confirm that most of these WNT genes were activated in R-femurs in presence of Scl-Ab (Fig. 2E). These strong correlations confirm the network maps generated by miRTargetLink 2.0 and indicated an important role of miR-183-5p in negatively regulating not only the WNT pathway, but also pathways involved in bone accrual.

**Figure 2.**
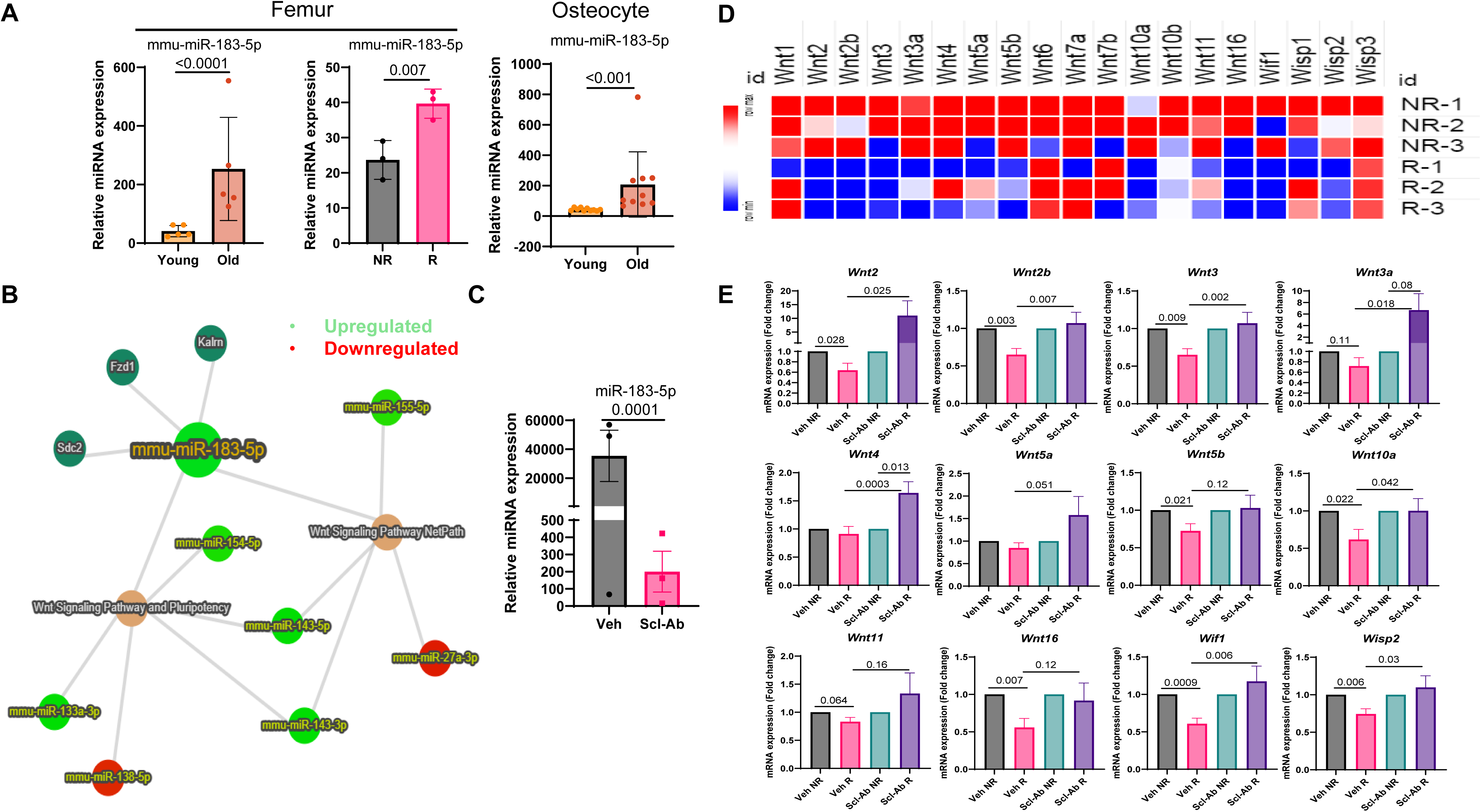
WNT pathway gene regulation by miR-183-5p during skeletal aging. (A) Differentially up-regulated miR-183-5p in O- vs Y- osteocyte enriched population, R- vs NR-femurs and O- vs Y- femurs. The statistical comparison was done using a two-tailed unpaired t-test. (B) The network analysis for miR-183-5p was done using miRTargetLink 2.0. (C) Heat map of WNT pathway genes in R-femurs vs NR-femurs. (D) mRNA from vehicle- or Scl-Ab-treated R-and NR-femurs from cumulative 16 Gy (8Gy on days 1 and 3) focally radiated mice, were collected on day 14 post radiation. WNT pathway genes were analyzed by qRT-PCR. Data was analyzed using a two-way ANOVA, with a post-hoc test: Dunnett’s test.

### Regulation of the pro-inflammatory signature by miR-155-5p during skeletal aging

Out of the 5 miRs that had shared upregulation between R-bones and O-osteocytes, miR-155-5p showed correlations with an important hallmark of skeletal aging, the proinflammatory SASP (Fig. 3A). Interestingly, miRTargetLink 2.0 analysis revealed that miR-155-5p regulated the TNF-alpha NF-kB signaling pathway, the chemokine signaling pathway, and the WNT signaling pathway (Fig. 3A), together with miR-183-5p as discussed above. To visualize the miR sequencing data analysis as a graph, significant upregulation was observed in the raw data reads for miR-155-5p in R-femur and O-osteocyte skeletal aging conditions, while a strong trend was seen in O-femurs (Fig. 3B). Furthermore, miR-seq data from Scl-Ab-treated mice showed lower expression of miR-155-5p, as compared to vehicle-treated mice (Fig. 3C). To further understand the correlation between miR-155-5p, the SASP and WNT pathway, we performed qRT-PCR on vehicle and Scl-Ab treated NR- and R- femurs. Surprisingly, Scl-Ab not only suppressed several SASP genes, such as *Ccl2*, *Ccl7*, *Cxcl2*, *Il18*, *Icam1* and *Mmp12*, but also suppressed senescence markers *Cdkn1a* (*p21*) and *Cdkn2a* (*p16^Ink4a^*) (Fig. 3 D, E).

**Figure 3.**
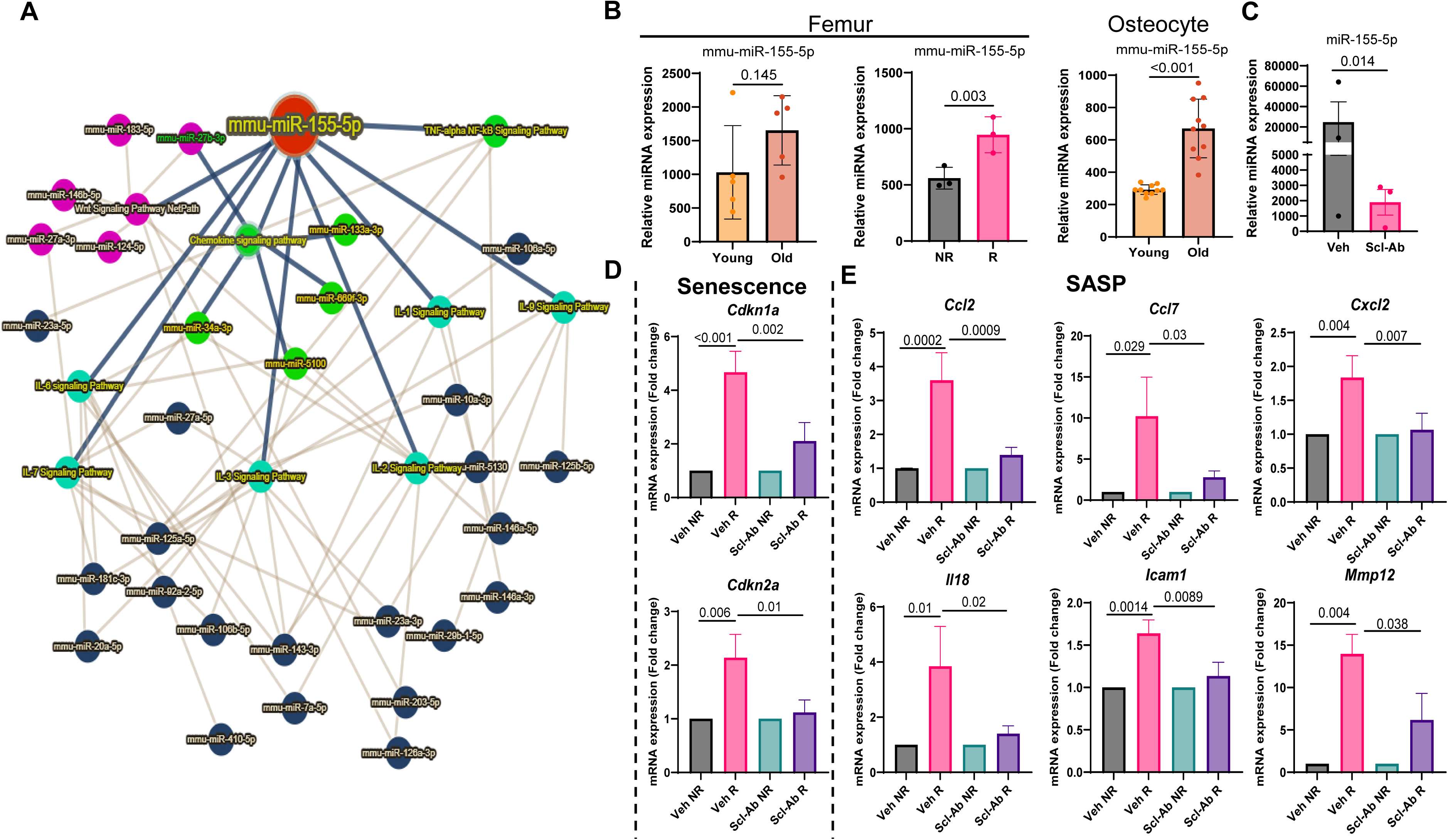
Pro-inflammatory SASP gene regulation by miR-155-5p and Scl-Ab during skeletal aging. (A) The network analysis for miR-155-5p was done using miRTargetLink 2.0. (B) Differentially up-regulated miR-155-5p during O- vs Y- osteocyte enriched population, R- vs NR-femurs and O-vs Y- femurs. The statistical comparison was done using a two-tailed unpaired t-test. (C) Gene expression analysis for senescence markers *Cdkn1a* (*p21*) and *Cdkn2a* (*p16^Ink4a^*) in vehicle- or Scl-Ab-treated R-and NR-femurs. Data was analyzed using 2-way ANOVA, followed by Dunnett test. (D) Gene expression analysis for several SASP markers in vehicle- or Scl-Ab-treated R-and NR-femurs. Data was analyzed using paired t-test for NR and R femurs from same animals and using 2-way ANOVA, followed by Dunnett test, for comparison between vehicle- or Scl-Ab-treated R-and NR-femurs.

### Regulation of FoxO1 by miR-135a-5p and WNT signaling

Among the two common downregulated miRs, miR-135a-5p correlated with the upregulation of some mRNA targets from our previously published mRNA seq data^12^. When raw values of miRNA expression were compared, miR-135a-5p levels are significantly reduced in O-femurs, R-femurs, and O-osteocytes (Fig. 4A). Among the gene targets of miR-135a-5p using miRTargetLink 2.0 analysis, *Foxo1* is identified as a prominent predicted target (Fig.4B). The schematic of the mouse *Foxo1* transcript (ENSMUST00000053764.5) shows a miR-135a-5p binding site within the *Foxo1* 3′UTR (Fig. 4C). It has been shown that elevation of *Foxo1* levels can suppress WNT signaling pathway^21^. So, we next tested, if WNT activation by Scl-Ab will regulate *Foxo1* mRNA levels. Indeed, qRT-PCR data shows that *Foxo1* was significantly elevated in R-femurs, confirming the inverse correlation between miR-135a-5p and *Foxo1* (Fig. 4D). Furthermore, elevated WNT activation by Scl-Ab treated R-femurs had reduced levels of *Foxo1* mRNA levels (Fig. 4D), confirming previously established relationship between WNT signaling pathway and FOXO1.

**Figure 4.**
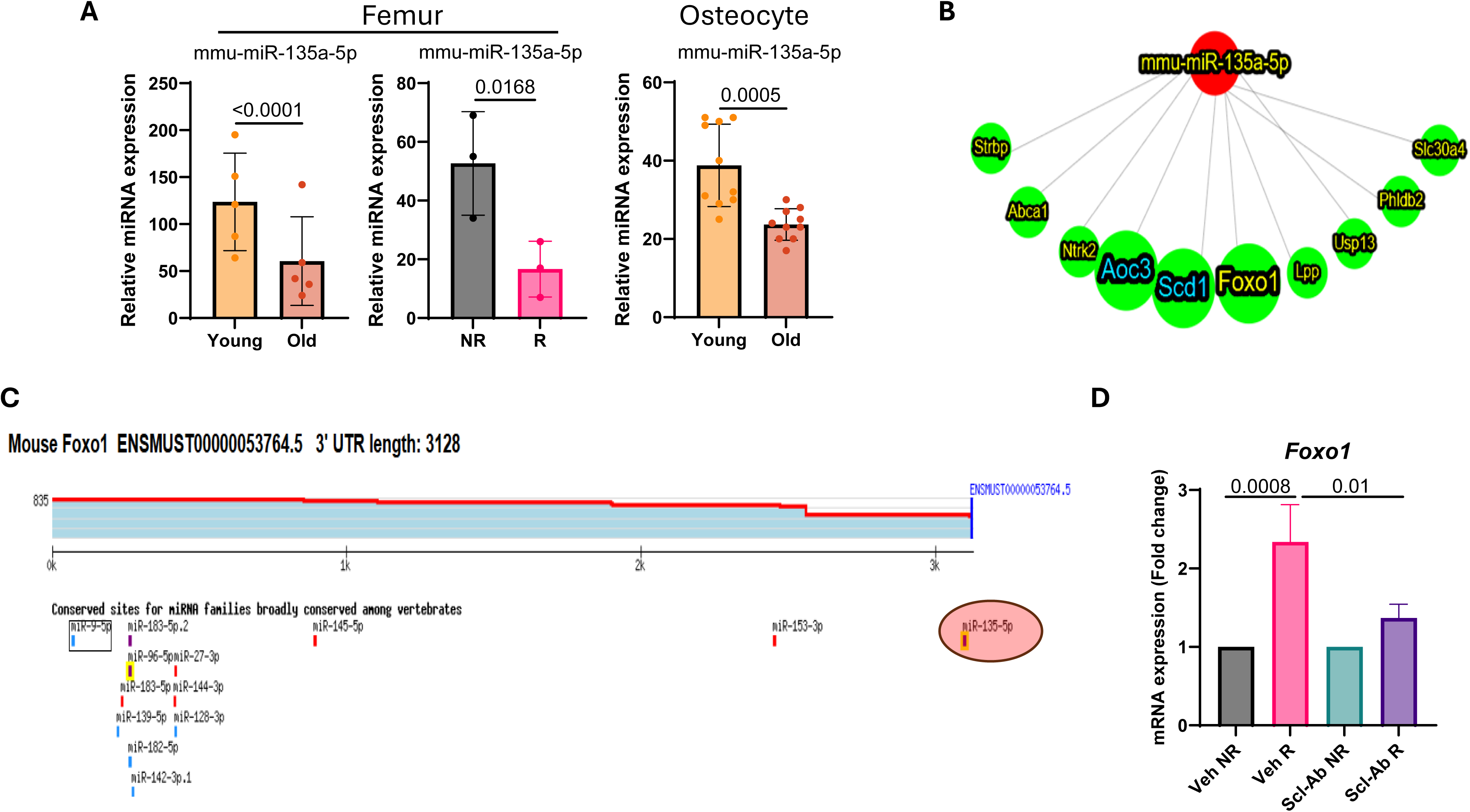
Regulation of Foxo1 by miR-135a-5p and Scl-Ab during skeletal aging. (A) Differentially down-regulated miR-135a-5p in O- vs Y- osteocyte enriched populations, R- vs NR-femurs and O- vs Y- femurs. (B) The network analysis for miR-135a-5p was done using miRTargetLink 2.0, identifying *Foxo1* as a key target. (C) Mouse target scan analysis identifying miR-135a-5p in the 3’UTR sequence of *Foxo1*. (D) Gene expression analysis for *Foxo1* in vehicle-or Scl-Ab-treated R-and NR-femurs. Data was analyzed using 2-way ANOVA, followed by Dunnett’s test.

### miRNAs that regulate Bone Marrow Adiposity and Bone homeostasis

Previously, using RNA-seq data, we have shown that R-femurs had high expression levels of genes related to BMAd^12^. Here, miRNA-seq data from Scl-Ab- and Veh-treated radiated femurs revealed that miRs that regulated BMAd-related genes, such as *Rxra* (miR-7b-5p) , *Scd1* (miR-7b-5p, miR-223-3p and miR-135a-5p), and *Aoc3* (miR-135a-5p), showed a trend to increase in Scl-Ab-treated R-femurs as compared to the vehicle-treated R-femurs (Supp. Fig.4).

When we compare the significantly expressed miRs that were down-regulated in Scl-Ab-treated R-femurs as compared to vehicle-treated up-regulated miRs, we identified 9 common miRs (Fig. 5A). Among these 9 common miRs, we focus here on miR-133a-3p specifically, since miRTargetLink 2.0 analysis revealed that miR-133a-3p negatively regulates several nodes and pathways that link to bone homeostasis, such as BMP signaling, WNT signaling, skeletal development and bone mineralization (Fig. 5B). Interestingly, miR-133a-3p downregulates genes involved in bone formation, *Runx2* and *Igf1r*, confirmed by our mRNA seq data^12^. As shown in our data, upregulation of miR-133a-3p in our cumulative 16Gy R-femurs at 2 weeks was reversed significantly by Scl-Ab treatment in R-femurs (Fig. 5C). Interestingly, there was a non-significant strong trend in upregulation of miR-133a-3p in miRNA-seq data from our 16Gy R-femur samples at 2 weeks but was significantly upregulated in both O-femurs and O-osteocyte samples and was among the 13 common miRs in these samples (Fig.1B).

**Figure 5.**
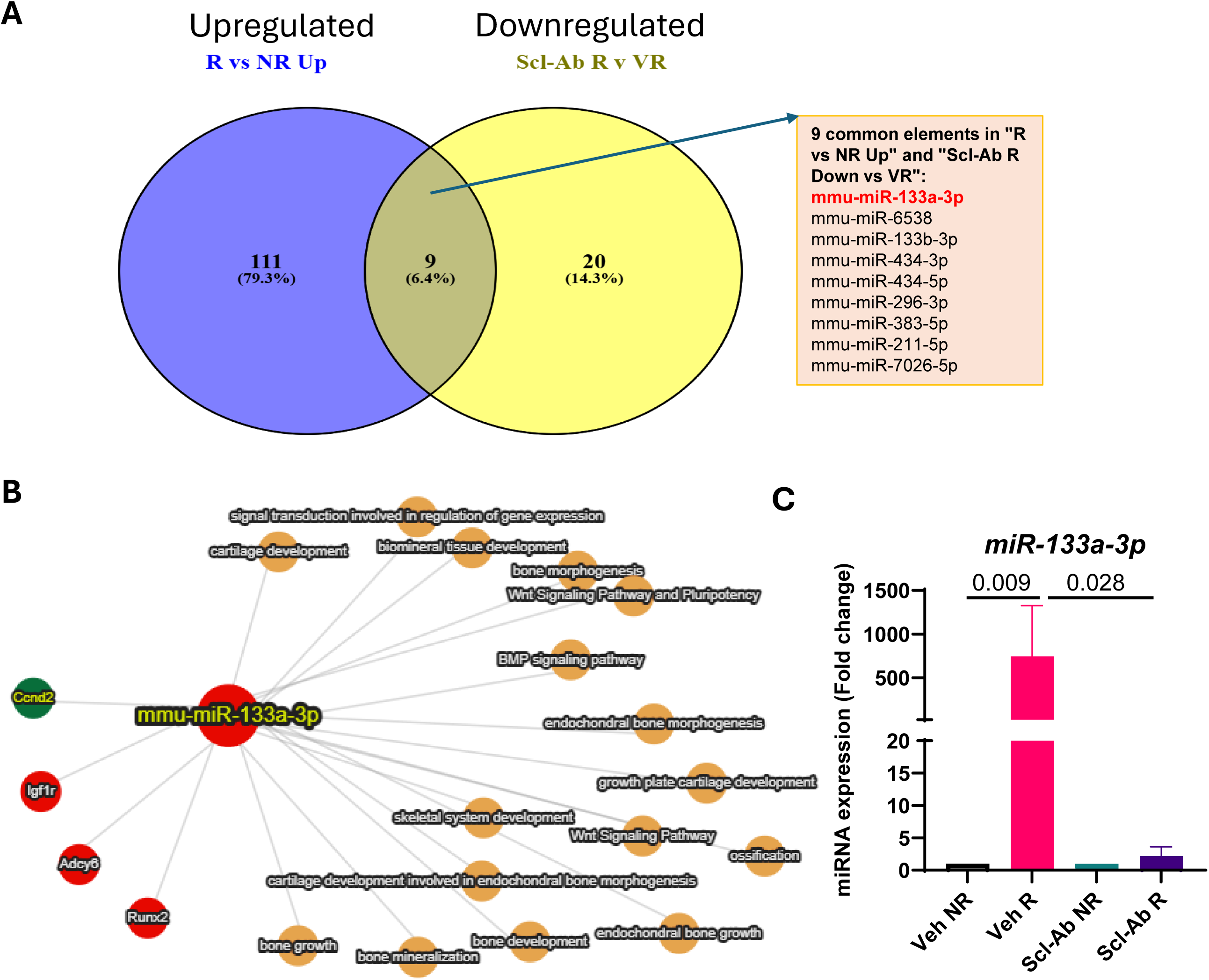
Role of miR-133a-3p in skeletal aging. (A) Venn diagram shows common or exclusive up-regulated miRs in R- vs NR-femurs and common down-regulated miRs in Scl-Ab-R vs Veh-R. (B) The network analysis for miR-133a-3p was done using miRTargetLink 2.0, identifying bone development pathways as key target pathways. (C) Raw data analysis of miR-133a-3p in vehicle- or Scl-Ab-treated R-and NR-femurs. Data was analyzed using 2-way ANOVA, followed by Šídák’s multiple comparisons test.

## Discussion

Explanations for the changes in the molecular signatures that regulate skeletal aging are still being explored. In this study, we identified miR-based molecular targets to several important pathways that regulate skeletal aging, including WNT signaling pathway, senescence and SASP and BMAd-related change. It was not until recently that mechanisms were more clearly defined in both pathological and physiological skeletal aging^10–12^. Using miRNA-seq data from three skeletal aging conditions, O-femurs, O-osteocytes and R-femurs, and the miRNA-seq data from Scl-Ab treated animals, we were able to identify miRs that control key pathways of skeletal aging and their regulation by WNT pathway agonist Scl-Ab (Fig.6).

**Figure 6.**
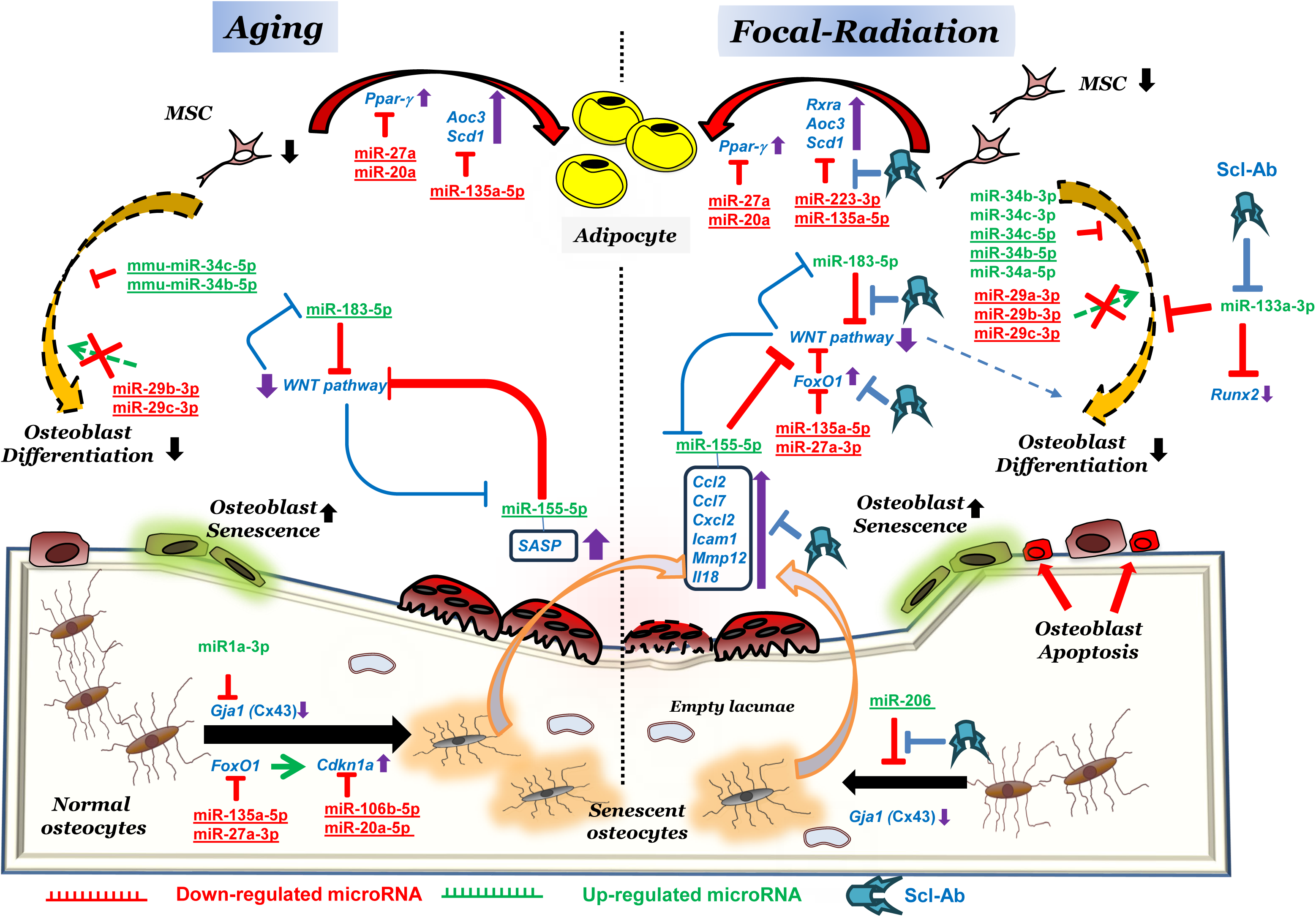
Schematic of miRNAs regulating skeletal aging. The figure depicts the important phenotypic changes that determine the cell fates seen during aging or focal radiation with: (i) reduced mesenchymal stem cell (MSC) numbers, (ii) increase in adiposity, (iii) reduction in osteoblast differentiation, (iv) reduction in mineralization, (v) increase in cellular senescence of bone environment, and (vi) reduced cellular communication because of osteocyte cell death, senescence and canalicular shrinking. Molecular changes such as microRNAs (miRs)-based regulation of gene expression have direct correlations with increased adiposity, reduced bone formation and increased senescence. The up-regulated miRs are represented in green and are shown to either target a gene (in blue) or a process. Similarly, the significantly downregulated miRs (in red) have been shown by us to correlate with increased expression of their corresponding genes either promote a process or inhibit one. Common miRs between aging and focal radiation are underlined.

Several miRs were differentially expressed in one or more of the three different skeletal aging conditions but were not expressed among all conditions. Figure 6 summarizes the functional relationships among these miRs and captures their unique or shared regulation between physiological and accelerated skeletal aging. These miRs and the pathways they control are strongly validated as shown by the miRTargetLink 2.0 analysis and from the literature survey done during the writing of this manuscript.

Our initial search for a common pathway that regulates skeletal aging led us to miR-183-5p, which was upregulated among the three skeletal aging conditions, while miR-671-5p and miR-135a-5p were the two common downregulated miRs. While miR-183-5p and miR-135a-5p had definite targets in bone and aging-related pathways, as shown by our data, we could not find known skeletal-related pathways for miR-671-5p. None of the genes upregulated in our previously reported mRNA-seq data^12^ seem to be regulated by miR-671-5p as shown by miRTargetLink 2.0 analysis (Supp. Fig. 5A). However, there is some indirect evidence that miR-671-5p regulates pathways linked to SASP/inflammation^22–25^ and WNT signaling^26^. On the other hand, there were some direct indications with miRTargetLink 2.0 that miR-183-5p regulates the WNT signaling pathway^27^, which we could validate using qRT-PCR data.

One of the key hallmarks of skeletal aging is cellular senescence with markers such as *Cdkn1a* (*p21*) and *Cdkn2a* (*p16^Ink4a^*), and miR106b^28,29^, miR20a^30^ and miR19b-3p^31^ (which have been shown to down-regulate p21) were significantly suppressed in both R-femurs and O-osteocytes. miR-17-3p, which is specifically down-regulated in R-bones, is also known to regulate p21 expression^32,33^. miR-106b, miR-20a, miR-17 and miR-19b, which were shown to be downregulated in several models of aging^32^, were also down-regulated in the R-bones, and correlated with the increase in transcript levels of p21 post-radiation. This correlation putatively supports the similarity between senescence mechanisms during radiation and aging.

FoxO1 is a transcription factor upstream of p21 and p27^34,35^ , regulated by miR-135a-5p^36,37^, and may also be a marker of senescence^38^. miR-27a-3p is up-regulated in liver cancer cells and adipose tissue with downregulation of *Foxo1*^39^. Similarly, we found that miR-27a-3p was significantly down-regulated in both R-femurs and O-osteocytes (Fig.1A), correlating with the upregulation of *Foxo1* transcript. miR-27a-3p also suppresses *Pparγ* (Supp. Fig. 5C) and may mediate the inverse relationship between adipogenesis and osteogenesis ^40^. We have previously reported increased adipocyte accumulation in R-bones at the expense of a limited number of MSCs, leading to reduced bone formation rate^13^. Similarly, levels of *Rxra* (regulated by miR-7b-5p), *Scd1* (regulated by miR-7b-5p, miR-223-3p and miR-135a-5p), and *Aoc3* (regulated by miR-135a-5p), are responsive to Scl-Ab, supporting an important role of WNT pathway in regulating BMAd-related genes. Another hallmark of senescence is the pro-inflammatory SASP, and identification of miR-155-5p that regulates SASP as a key upregulated miR in our skeletal aging samples is consistent with miR-155-5p also being an miR biomarker for skeletal aging^41,42^. Interestingly, the key miRs that appear to regulate skeletal aging also target WNT pathways.

Studies have attempted to identify miRs as biomarkers that could predict aging and osteoporosis and in some instances bone quality. For example, miR-20a-5p, which was among those miRs in-common between R-femurs and O-osteocytes (Fig.1A) and a known promoter of osteogenesis^43^, could potentially mediate low bone formation upon its downregulation. Another study found a direct correlation between miR-29b-3p with improved bone formation rate/bone surface^44^. miR-29a appears to be an enhancer of mineral deposition^45^, while miR-29b may promote osteoblast differentiation^46^. All three miRs of the miR29 cluster (miR-29a-3p, miR-29b-3p and miR-29c-3p) were significantly downregulated in the R-bones and O-osteocytes (Fig. 1A). In the aged osteocyte samples, miR-29b-3p and miR-29c-3p were significantly downregulated, but not miR-29a-3p. A conundrum here as that miR-29 is also known to induce senescence in certain models^47,48^, but likely not in bone, since in both aged osteocytes as well as R- bones the miR-29 cluster was reduced in expression, and correlated with an increase in markers of senescence. We did not see any significant changes in our miRNA-seq data from Scl-Ab treated mice, suggesting that WNT signaling may not regulate miR-29 cluster. One of the keys miRs that negatively regulates bone homeostasis and osteogenesis is miR-133a-3p^49–51^. Our data show that R-femurs had upregulated miR-133a-3p as compared to NR-femurs, while this increase was nullified in the presence of Scl-Ab- treated mice. This indicates that miR-133a-3p is a key miR, one that could help to define at least accelerated skeletal aging secondary to irradiation.

Conversely, the miR-34 cluster (miR-34b-3p, miR-34c-3p, miR-34c-5p, miR-34b-5p and miR-34a-5p) which is linked to suppression of osteoblast differentiation and reduced bone accrual by inhibiting the *Satb2* gene^52^, was significantly elevated in the R-bones, and in O-femurs (miR-34c-5p and miR-34b-5p). miR-1a-3radiated bones and miR-206 down-regulate (*Gja1*) connexin-43^53,54^ (Supp. Fig.5B) and may be linked with the reduced osteocytic canalicular connections as reported in aged-^55^ and radiated-bones^13^. Interestingly, miR-1a-3p was significantly elevated in O-osteocytes and O-femurs.

Finally, we would like to address the possibility of alternative mechanisms for radiotherapy-induced bone damage (i.e., other than direct DNA damage). We have previously demonstrated that osteoblast apoptosis and MSC fate switching to adipogenesis were significant contributors for radiotherapy-related bone deterioration^13,14^. Apart from the direct effects of radiation on the DNA double strand, epigenetic regulators like EZH2 seem to suppress osteoblastogenesis upon exposure to radiation^56^. Furthermore, long term sustained epigenetic changes are observed in bone marrow hematopoietic progenitor and stem cells when exposed to low dose ionizing radiation^57^. EZH2 is also regulated by miR, and our sequencing data confirmed that both miR-101 and miR-126, which are shown to target EZH2^58,59^, were significantly downregulated in both R-femurs and O-osteocytes (Fig. 1A). Epigenetic regulation post-clinical radiotherapy is multifaceted and needs to be further investigated.

Here we have provided evidence that specific miRs have shared roles among models of skeletal aging and may also regulate pathways in common between physiological and accelerated skeletal aging. Using *in silico*, *in vitro* and *in vivo* methods, our study identified and validated key miRs and their respective mRNAs that regulate either the WNT pathway, senescence or SASP markers and some other processes in bone remodeling (osteoblast differentiation, osteocyte maintenance and adipocyte differentiation). Identification of these miR-mRNA correlations allows an expansion in the current knowledge about the molecular pathways regulating physiological and pathological osteoporosis.

## Author Contributions

**Achudhan David:** Experimentation and Data collection and analysis; writing – review and editing. **David G. Monroe:** Data collection and analysis; funding acquisition; writing – review and editing. **Ling Qin:** funding acquisition; writing – review and editing. **Sundeep Khosla:** funding acquisition, writing – review and editing. **Robert Pignolo:** Conceptualization; project administration; resources; writing – original draft; writing – review and editing; **Abhishek Chandra:** Conceptualization; project administration; funding acquisition; resources; data analysis; writing – original draft; writing – review and editing.

## Acknowledgements

This work was made possible by the Robert and Arlene Kogod Professorship (to RJP), R01 AG082681 (A.C), P01 AG062413 (S.K., D.G.M.), R01 AG086085 (S.K., D.G.M.), R01 AG076515 (S.K., D.G.M.), Hevolution HR-GRO-23-1199144-8 (S.K., D.G.M.), NIH/NIAMS R01AR066098 (L.Q.) and by the Center for Clinical and Translation Science (to D.M. and A.C.).

