## Supplemental figures for "MicroRNA Networks Driving Skeletal Aging and WNT Pathway Modulation"

Supplementary Figure.1

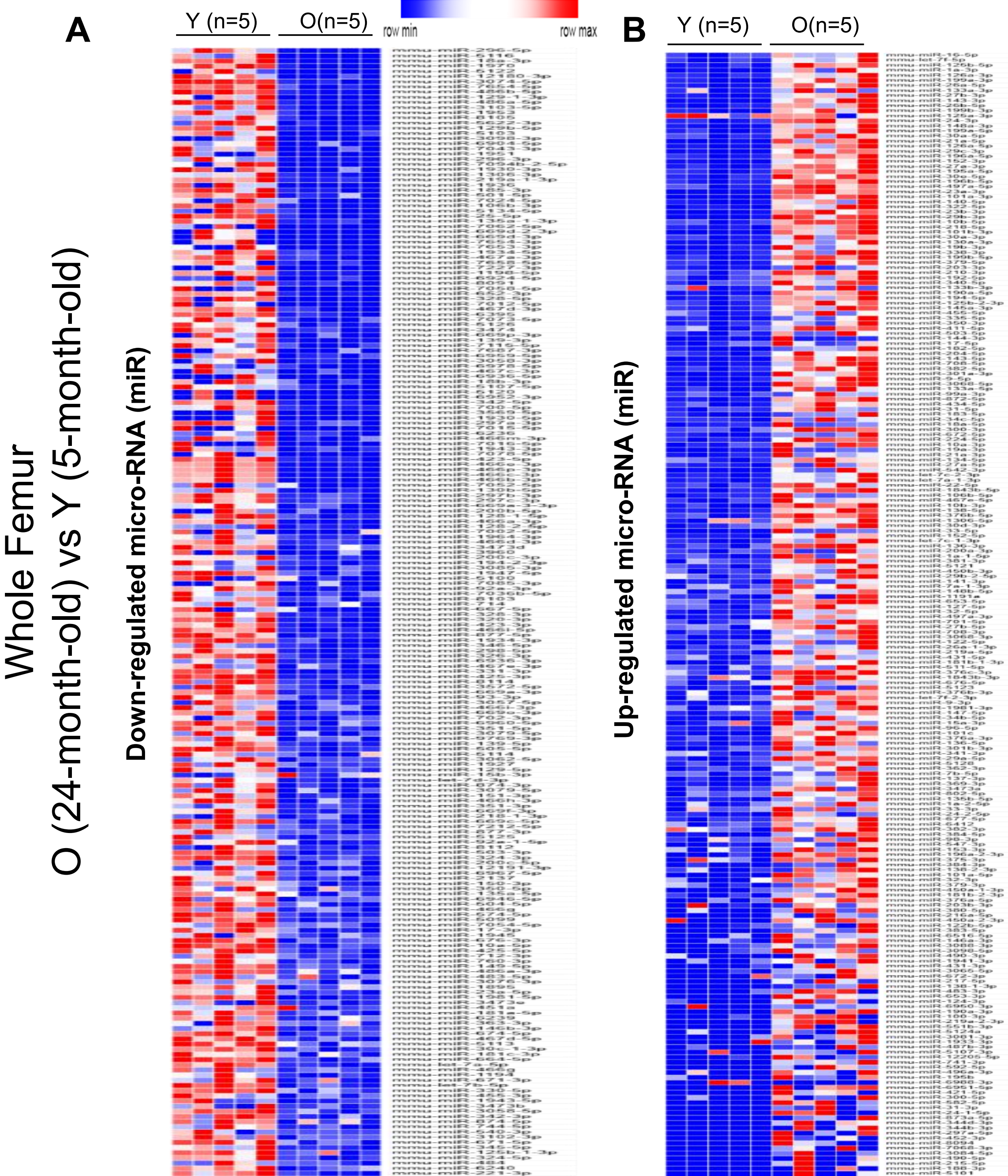

Supplementary Figure.2

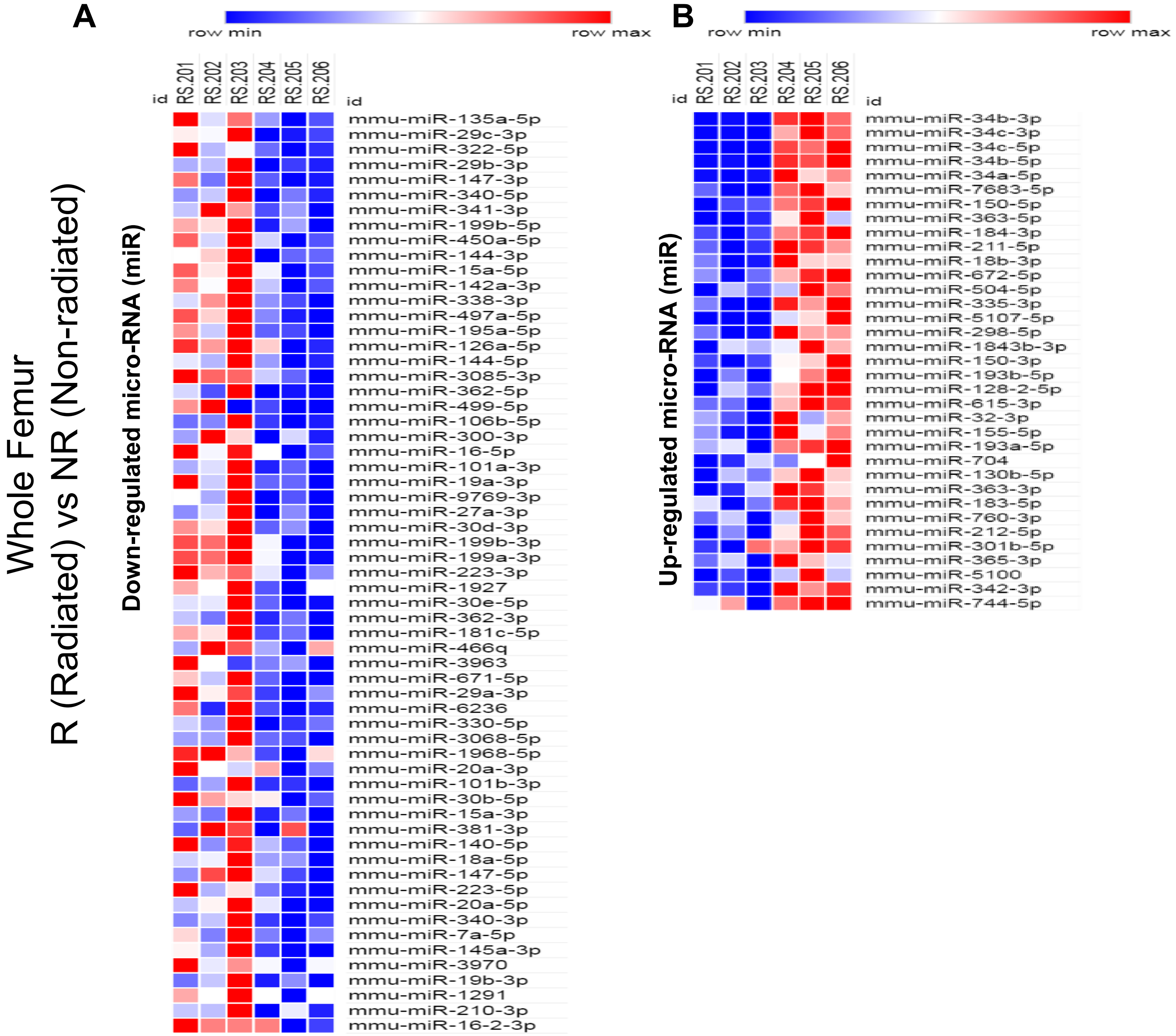

Supplementary Figure.3

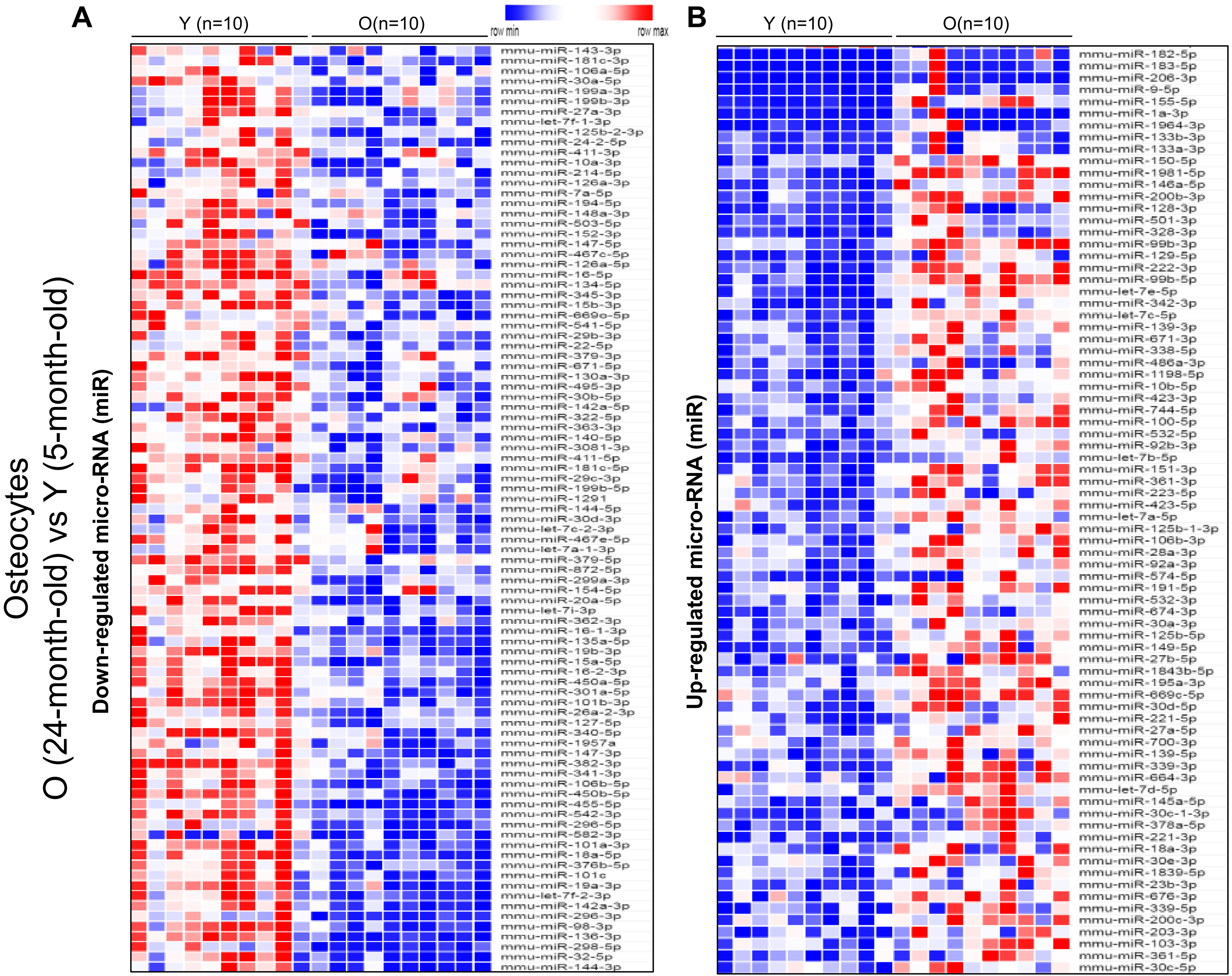

Supplementary Figure.4

A

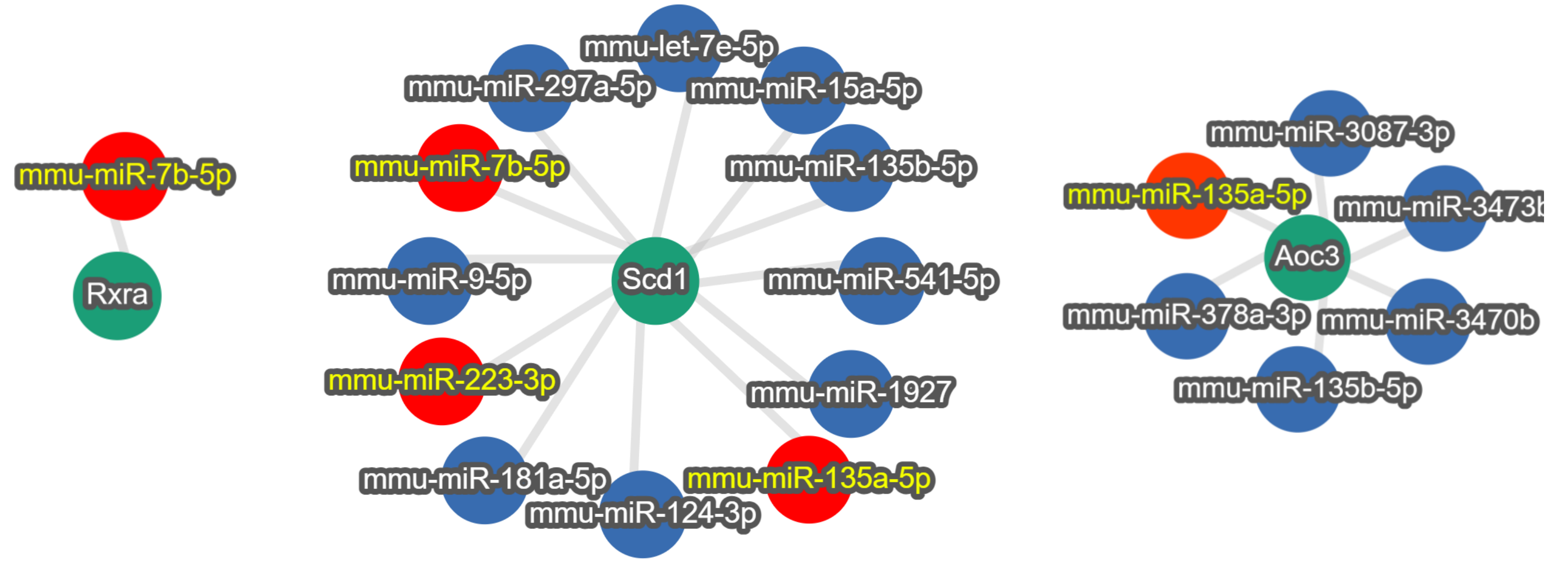


B

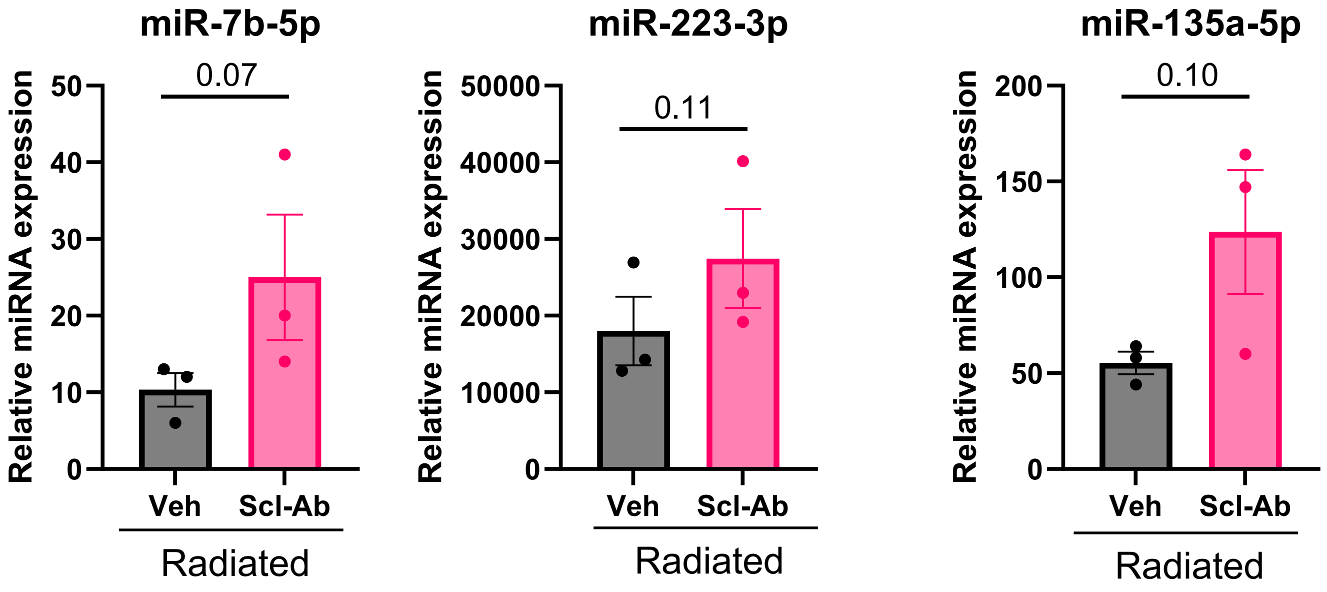

Supplementary Figure.5

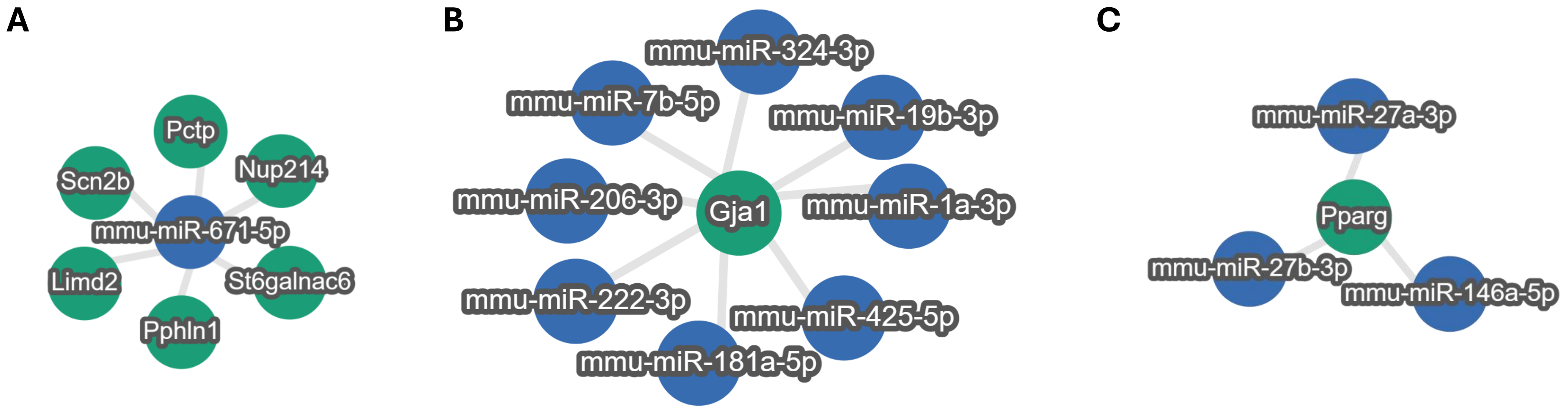

**Supplementary Figure Legend**

**Supplementary Figure 1. Differentially regulated miRNAs in aged and young femur**

mRNA from femurs from 5-month- and 24-month- old mice were collected and miRNA-sequencing was performed. Differentially expressed miRs are plotted as heat maps of commonly down-regulated miRs (A) and up-regulated miRs (B).

**Supplementary Figure 2. Differentially regulated miRNAs in radiated femur**

mRNA from femurs from 16Gy radiated (R) and non-radiated (NR) femurs 4-month- old mice, where femoral metaphysis was collected on day 14 post-radiation and miRNA-sequencing was performed. Differentially expressed miRs are plotted as heat maps of commonly down-regulated miRs (A) and up-regulated miRs (B).

**Supplementary Figure 3. Differentially regulated miRNAs in aged and young osteocytes**

mRNA from enriched osteocytes from femurs of 5-month- and 24-month- old mice were collected and miRNA-sequencing was performed. Differentially expressed miRs are plotted as heat maps of commonly down-regulated miRs (A) and up-regulated miRs (B).

**Supplementary Figure 4. BMAd-related miRNAs**

(A) The network analysis for BMAd-related genes *Rxra* , *Scd1* and *Aoc3* were done using miRTargetLink 2.0, identifying miRs that regulate these genes. (B) Raw data analysis of miR-7b-5p, miR-223-3p, and miR-135a-5p in vehicle- or Scl-Ab-treated mouse femurs. The statistical comparison was done using a two-tailed unpaired t-test.

**Supplementary Figure 5. Miscellaneous mRNA and miRNA correlation regulating skeletal aging**

(A) The network analysis for miR-671-5p using miRTargetLink 2.0. miR-671-5p is one of the two miR that is commonly down-regulated in O- vs Y- osteocyte enriched population, Rvs NR-femurs and O- vs Y- femurs. (B) The network analysis for *Gja1* using miRTargetLink 2.0. (C) The network analysis for *Pparg* using miRTargetLink 2.0.
